# Synaptic dysfunction and dysregulation of extracellular matrix-related genes in dopaminergic neurons derived from Parkinson’s disease sporadic patients and with *GBA1* mutations

**DOI:** 10.1101/2023.04.10.536143

**Authors:** Idan Rosh, Utkarsh Tripathi, Yara Hussein, Wote Amelo Rike, Andreea Manole, Diogo Cordeiro, Henry Houlden, Jurgen Winkler, Fred Gage, Shani Stern

**Author notes:** Equally contributed (Idan Rosh, Utkarsh Tripathi).

## Abstract

Parkinson’s disease (PD) is a neurodegenerative disease with both genetic and sporadic origins. In this study, we investigated the electrophysiological properties, synaptic activity, and gene expression differences in dopaminergic (DA) neurons derived from induced pluripotent stem cells (iPSCs) of healthy controls, sporadic PD (sPD) patients, and PD patients with GBA1 mutations. Our results demonstrate reduced sodium currents and synaptic activity in DA neurons derived from PD patients with GBA1 mutations, suggesting a potential contribution to PD pathophysiology. We also observed distinct electrophysiological alterations in sPD DA neurons that were dependent on the age of disease onset. RNA sequencing analysis revealed unique dysregulated pathways in early and late-onset sPD neurons, further supporting the notion that molecular mechanisms driving PD may be different between PD patients. In agreement with our previous reports, ECM and focal adhesion genes were the top dysregulated pathways in DA neurons from sPD patients and from patients with GBA1 mutations. Overall, this study gives further confirmation that the convergent functional phenotypes of DA neurons derived from PD patients are synaptic abnormalities and at the transcriptome level, ECM and focal adhesion pathways are highly involved in PD pathology across multiple PD-associated mutations as well as sPD.

## Introduction

Parkinson’s disease (PD) is a neurodegenerative disease characterized by progressive extrapyramidal motor dysfunction (1) and is considered to be an age-related disease (2-5). Compared to other neurological diseases, its prevalence has almost doubled in the past 25 years (6). Given asymptomatic people who remain undiagnosed due to the disease’s slow progression and subsequent late onset of symptoms, the incidence of PD is assumed to be considerably greater than what is officially reported (6-8). The development of PD’s motoric deficits is associated with a progressive loss of dopaminergic neurons in the substantia nigra pars compacta and the subsequent depletion of dopamine levels in the striatum (9). Albeit predominantly dependent on the presence of motor deficits such as bradykinesia, rigidity, and tremor for its diagnosis, PD is characterized by a range of non-motoric preclinical features such as rapid eye movement sleep behavior disorder, anxiety disorders, anemia, depression, and constipation. Most interestingly, the mentioned non-motoric features, which are associated with abnormalities of the serotonergic, noradrenergic, and cholinergic systems, can, in some cases, manifest years before the disease onset and are linked to pathology in various parts of the nervous system (10-12). With that said, many aspects of PD’s etiology are still considered enigmatic, even two centuries after its initial description by Dr. James Parkinson (13). Disease-modifying agents for PD are an unmet requirement; it is still possible to alleviate symptoms with multiple types of medication, thus allowing PD patients to maintain their active life for several years longer (14, 15).

The majority of PD cases, about 85%, are considered a non-genetic disorder - a ‘sporadic’ origin. Around 15% of PD cases are known to be inheritable and attributable to defects in distinct sets of genes in a specific genetic locus and are considered familial PD’ (16-19). Exhibiting an overlapping presentation, neuropathology, and disease mechanism, this monogenic form of the disease is distinct by its varied degree of increased risk for PD and noteworthy early age onset; some have a slow progression of motoric and mental symptoms, according to the distinct carried risk factors (20, 21). In contrast, sPD is mainly characterized by a late-age onset, and the patients usually manifest motoric symptoms after the age of 60-65. The minority of sPD patients present parkinsonian symptoms before 40-50 years of age and are classified as cases of early-onset. Apart from age onset, the mentioned forms of sPD differ by distinct disease progression, clinical features, and response to medication (19). Identifying the basis for the differential disease characteristics between early and late-onset sPD may be crucial for understanding the underlying mechanisms of PD pathogenesis and for the discovery of novel therapeutic targets (22).

Several mechanisms have been suggested to be involved in the etiology of sPD, including abnormal handling of misfolded proteins by the proteasomal enzymatic activities, dysfunction in the mitochondrial electron transport chain, increased oxidative stress, and other pathogenic dysfunctions (23, 24). Pathologically, sPD is often characterized by the accumulation of α-synuclein in Lewy bodies and the degeneration of DA neurons in the substantia nigra pars compacta (25). Recent studies on sPD cases have demonstrated alterations in synaptic activity and dysregulated extracellular matrix pathways in midbrain neurons derived from PD patients (26, 27).

The motor symptoms of sPD appear when the striatum’s dopamine concentration falls by roughly one-third and more than half of the patient’s substantia nigra pars compacta DA neurons are lost, implying a possible involvement of a compensatory mechanism in the early stages of the disease (28-30). However, most sPD patients lack a definitive genetic basis making it difficult to create experimental models and find well-targeted medicines for disease management. Despite the lack of a clear genetic basis, numerous single nucleotide variants have been identified as contributing risk factors for sPD, suggesting that the condition might stem from a correlative composition of causative genetic factors (31-35). Studies indicate that 5–15% of sPD patients were found to be carrying β-glucocerebrosidase (*GBA1*) gene mutations, making it the most prevalent genetic risk factor for sPD to be identified (36, 37). With the exception of an earlier age of onset, rapid progression of motor symptoms, and higher cognitive dysfunction, GBA-PD is not clinically distinct from sPD (36, 38-41). Furthermore, both conditions involve identical pathology, including nigrostriatal dopamine loss and the deposits of aggregated α-synuclein in the form of Lewy bodies (42-44).

*GBA1* gene encodes glucocerebrosidase (GCase), a lysosomal enzyme responsible for the cleavage of glucose from glucosylceramide (GlcCer) and glucosyl sphingosine (GlcSph) (45). Mutations in both alleles of *GBA1* results in Gaucher’s disease (GD), an autosomal recessively inherited lysosomal storage disease (46), while its heterozygous loss-of-function mutations are one of the most common genetic risk factors identified in PD (36, 47, 48). The *GBA1* gene mutations have also been found to be significantly more common in PD patients than in non-affected individuals in over 50 population studies that have looked at the *GBA1* gene in PD patients (49). However, fewer *GBA1* mutations have been found in patients with PD compared to those with GD (49). Between *GBA1* mutations, two-point mutations, N370S (p.N409S) and L444P (p.L483P), are the most common GD-associated ones (45, 50). These mutations vary across different populations, with N370S being the most common mutation among Ashkenazi Jews (AJ), accounting for about 70% of mutant alleles, and the L444P among the non-AJ European ancestors (51, 52). The E326K mutation in the GBA1 gene is a specific point mutation and non-pathogenic polymorphism that has been implicated in an increased risk for Parkinson’s disease and other synucleinopathies. Mutations in the *GBA1* gene, including the E326K variant, have been associated with reduced enzymatic activity and the subsequent accumulation of glycolipids in cells, leading to cellular dysfunction and neurodegeneration (36, 52, 53).

The frequency of *GBA1* mutation carriers is also high, 10–31%, among the European Ashkenazi Jewish population compared to the European non-Ashkenazi Jewish population, which is 2.9– 12% (36). The discovery of the association between the *GBA1* mutations and the increased PD risk has resulted in essential considerations and discoveries on PD’s pathogenesis(54, 55). Several methods have been employed to develop a mouse model of *GBA1*-linked Parkinson’s disease, such as knocking out *GBA1* or introducing point mutations, as well as chemically induced models (56). Although each approach provides distinct benefits, there is no single optimal mouse model for *GBA1*-PD, which underscores the importance of selecting the most suitable model. Unfortunately, there are no mouse models for sPD. Patients’ neurons derived from iPSCs hold good prospect in bridging this gap in seeking mechanisms of uncovering therapeutic approaches for PD neurodegeneration. This technology has greatly enhanced the molecular understanding of genetically complex diseases through better mirroring of its phenotypic manifestations by using live brain tissue models (57).

With the help of iPSC techniques, it has been shown that mutations in the *GBA1* gene result in disrupted mitochondrial function (58), increased endoplasmic reticulum (ER) stress (59), impaired lysosomal morphology and function (58, 60), decreased level of cathepsin D (61), enhanced aggregation of α-synuclein (62), and up regulation of monoamine oxidases (MAO) in DA neurons (63-65). Additionally, research using iPSC-derived DA neurons from PD patients with mutant *GBA1* demonstrated that these neurons had abnormalities in calcium homeostasis, an increase in α-synuclein and GlcCer levels, and a marked reduction in protein levels and GCase enzyme activity when compared to isogenic controls (62). Similar results were obtained from a twin study that looked at iPSC-derived DA neurons and discovered that *GBA1* mutations are connected to an elevated α-synuclein and impaired GCase activity (66). As reviewed by Tran et al. (57), though sPD accounts for a significant portion of all cases of PD, there are not many iPSC studies of sPD (26, 27, 67).

In this study, we concentrated on the neurophysiology and transcriptional alterations in DA neurons derived from PD patients with *GBA1* mutations and sPD patients. Our results revealed distinct and shared dysregulated pathways in both *GBA1*-associated and sPD cases, some that are shared with our previous reports for other mutations (26, 27, 67). In *GBA1*-associated PD, we found a significantly reduced influx of sodium currents and hypo excitability in DA neurons as well as dysregulation in ECM-receptor interactions, focal adhesion, and more pathways. For early and late-onset sPD DA neurons, we observed a dysregulation in pathways such as mitochondrial function, synapse, ECM, and focal adhesion. We further measured spontaneous synaptic activity and observed a significantly reduced rate of spontaneous synaptic activity and smaller amplitudes of synaptic events from DA neurons derived from *GBA1* PD patients compared to healthy individuals. In sPD the synaptic activity impairments depended on the disease onset age (increased for late-onset and decreased for early-onset, compared to healthy controls). Overall, this study strengthens our previous studies connecting synaptic impairments and dysregulation of ECM and focal adhesion-related genes to PD.

## Methods and Materials

### Midbrain DA differentiation

To generate *in vitro* midbrain DA neurons, we employed a protocol previously developed and reported in our earlier studies (26, 68, 69). We differentiated midbrain DA neurons from 3 control individuals, 3 PD patients with the E326K mutation in the GBA1 gene, and 4 sPD patients. Briefly, the human iPSCs were dissociated, replated, and allowed to proliferate before initiating differentiation at 50% confluency (day 0). A gradual transition from KSR medium (DMEM F-12 with Glutamax, KO-SR, NEAA, β-mercaptoethanol) to N2 medium (DMEM F-12 with Glutamax, N2 supplement) occurred from day 5 to day 10, followed by a switch to B27 medium (Neurobasal medium, B27 supplement, glutamax, BDNF, GDNF, TGFβ3, ascorbic acid, and Cyclic adenosine monophosphate (cAMP)) on day 11. Various small molecules were added during the differentiation, such as SB431542, LDN-193189, Smoothened agonist (SAG), and fibroblast growth factor (FGF) 8b. Neurons were dissociated and replated on days 20-25 and maintained in the B27 medium until day 30 when the base medium was replaced with Brainphys medium(70) to promote synaptic connections. Whole-cell patch clamp and RNA sequencing experiments were conducted on mature neurons (more than 7 weeks in differentiation).

### Immunocytochemistry

Coverslips with DA neurons were fixed in 4% paraformaldehyde (PFA) for 15 minutes at 37^0^ C. After washing with DPBS, and the cells were blocked and permeabilized in a solution of DPBS, 0.2% Molecular Grade Triton X-100, and 10% Donor Horse Serum for 1 hour. Primary antibodies were added to the blocking solution at 4°C overnight, with the following dilutions for DA neurons: TH (1:500) and MAP2 (1:500), and for hippocampal neurons: PROX1 (1:4000) and MAP2 (1:500). The next day, the coverslips were washed with DPBS and incubated with Alexa Fluor secondary antibodies, followed by counterstaining with DAPI staining solution (1:3000) for one hour at room temperature. The coverslips were rewashed, mounted on slides using Fluoromount-G mounting medium (0100-01, Southern Biotech), and allowed to dry overnight in the dark. The fluorescence signals were visualized using a Nikon A1-R confocal microscope, and images were processed using NIS elements 5.21 (Nikon) and microscopy image analysis software Imaris 9.8 (Oxford Instruments).

### RNA extraction, sequencing, and analyses

The total RNA was extracted from 3-5 million DA neurons per sample, derived from three patients with *GBA1* mutations, one patient with an early-onset sPD, one patient with a late-onset sPD, and three healthy control lines at 7-8 weeks post-differentiation, using the zymo RNA clean & concentrator kit as per the manufacturer’s instructions. The extracted RNA was reverse transcribed with the high-capacity cDNA synthesis kit from AB Biosystems.

To analyze the RNA sequencing data reads sequenced on a next-generation sequencing (NGS) platform were processed in FASTQ-format files. The sequences were trimmed using the Trimmomatic algorithm and quality-tested using FASTQC v0.11.8. Sequences were aligned to the hg38 human genome using STAR aligner v2.78a. Mapping was carried out using default parameters, filtering non-canonical introns, allowing up to 10 mismatches per read, and only keeping uniquely mapped reads. The expression levels of each gene were quantified by counting the number of reads that aligned to each exon or full-length transcript and normalized by its mean across all samples using HTseq v0.9.1. Differentially expressed genes (DEGs) were determined using DESeq1 v.2.11.40.7, and the p-value was adjusted for multiple hypotheses with the Benjamini-Hochberg procedure(71), which controls for the false discovery rate (FDR). Genes with an FDR > 0.05 were included in the analysis. A Gene Ontology (GO) enrichment test and KEGG pathway analysis were performed using expressAnalyst. A graphical representation of the protein network pathways analysis was constructed using nodes and edges, where each node represented a pathway connected by the edges (lines). The size of each node is proportional to the number of DEGs. An overrepresentation of GO terms and KEGG pathway was determined by FDR < 0.05.

### Patch-Clamp Recordings of Dopaminergic Neurons

Whole-cell patch-clamp recordings were performed from DA neuronal cultures. The recordings were conducted at 45-50 days of the differentiation. Culture coverslips were placed inside a recording chamber filled with HEPES-based artificial cerebrospinal fluid (ACSF) containing NaCl (139 mM), HEPES (10 mM), KCl (4 mM), CaCl2 (2 mM), D-glucose (10 mM), and MgCl2 (1 mM) at pH 7.4 and osmolarity adjusted to 310 mOsm at room temperature. Recording micropipettes with a tip resistance of 10-15 MΩ were filled with an internal solution containing K-gluconate (130 mM), KCl (6 mM), NaCl (4 mM), Na-HEPES (10 mM), K-EGTA (0.2 mM), GTP (0.3 mM), Mg-ATP (2 mM), cAMP (0.2 mM), D-glucose (10 mM), biocytin (0.15%), and rhodamine (0.06%) at a pH of 7.4 and osmolarity adjusted to 290-300 mOsm. Data were recorded in a sampling rate of 20 kHz using Clampex v11.1.

### Analysis of total evoked action potentials

To determine the total number of evoked action potentials, neurons were held in current clamp mode at a steady membrane potential of -60 mV with a constant holding current. Current injections were given in 3 pA steps over 400 ms, starting 12 pA below the steady-hold current required for a -60 mV membrane potential. The total number of evoked action potentials in response to the first 32 depolarization steps within the 400 ms recordings was counted and compared between the groups as a measure of the excitability.

### Input conductance and capacitance

The input conductance was determined by measuring the current while holding the cell in voltage-clamp mode at -70 mV and -50 mV, respectively, around the resting membrane potential. The input conductance was calculated by dividing the difference in currents by the difference in membrane potentials (20 mV). The capacitance was measured using the membrane test feature in the Clampex software.

### Analysis of sodium, fast, and slow potassium currents

Sodium and potassium current measurements were performed in voltage-clamp mode. Voltage steps of 400 ms in the range of -90 to 80 mV were produced while holding the cells at a voltage of -60 mV. The measured currents were normalized by the cell capacitance. The maximal outgoing current within a few milliseconds after a depolarization step was used to calculate the fast potassium current, while the current at the end of the 400 ms depolarization phase was used to calculate the slow potassium current. The amplitudes of the sodium and potassium currents were statistically analyzed to compare PD vs. control DA neurons at specific test potentials (-20 - 0 mV for the sodium current and 50 - 90 mV for the potassium current).

### Analysis of synaptic activity

Spontaneous excitatory post-synaptic currents (EPSCs) were recorded in a voltage-clamp mode to analyze synaptic activity. Neurons were held at -60 mV, and the amplitude and rate of synaptic activity were evaluated using a custom-written Matlab code. The cumulative distribution of EPSC amplitudes was calculated for each group. The EPSC event rates and amplitudes were calculated as the average of events per each recorded cell.

We additionally calculated the EPSC rate of large EPSCs whose amplitudes were greater than 30 pA. To analyze the synaptically active cells, neurons were considered synaptically active if they had at least three EPSCs during the 60 seconds of the EPSC recordings.

### Statistical analyses

Statistical analyses were performed using Clampfit-11.1 and Matlab software (R2021a, The MathWorks Inc., Natick, MA, 2000). Two-sample t-tests (two-tailed) were used to calculate p-values unless otherwise specified. A one-way ANOVA analysis was used to analyze the sodium and potassium current changes. All data values are presented as mean ± standard error (SE). A p-value of less than 0.05 was considered significant for all statistical tests.

## Results

Our study included ten iPSC lines, comprising three lines from healthy controls, three lines from PD patients with a *GBA1* mutation, and four lines from sPD patients. The early onset sPD group included three patients with an onset age of 46, 51, and 55, while the late onset sPD patient had a disease onset of 74 years. To investigate the intrinsic properties and synaptic activity of these cells, we differentiated the iPSCs into DA neurons (see Methods) and performed whole-cell patch clamp recordings (see Methods).

### *GBA1* mutant DA neurons exhibit reduced synaptic activity and a reduction in the sodium currents

Using a whole-cell patch-clamp, we recorded from a total of 53 control and 40 *GBA1* mutant neurons from 3 control individuals and 3 *GBA1* PD patients. Figure 1(a)-(b) show representative traces of evoked potentials in a control neuron and a *GBA1* mutant neuron, respectively. The total number of evoked potentials was significantly lower in *GBA1* mutant neurons compared to control neurons (Figure 1(c), p=0.04). Further, we compared the synaptic activity between control and *GBA1* mutant neurons measured by holding the neurons in a potential of -60mV. Figures 1(d)-(e) show examples recordings of EPSCs in control and *GBA1* mutant neurons, respectively. We observed a significant reduction in the mean amplitude of EPSCs in *GBA1* mutant neurons compared to control neurons (Figure 1(f), p=0.0053). Additionally, the EPSC rate was significantly lower in *GBA1* mutant neurons compared to control neurons (Figure 1(g), p=0.016). Figure 1(h) shows the cumulative distribution of EPSC amplitudes of *GBA1* mutant and control neurons. This distribution was left-shifted in the *GBA1* neurons compared to control neurons, indicating lower EPSC amplitudes. We observed no significant differences in the capacitance of *GBA1* compared to control neurons, indicating that the neurons’ total surface area was similar (Figure 1(i)).

**Figure 1:**
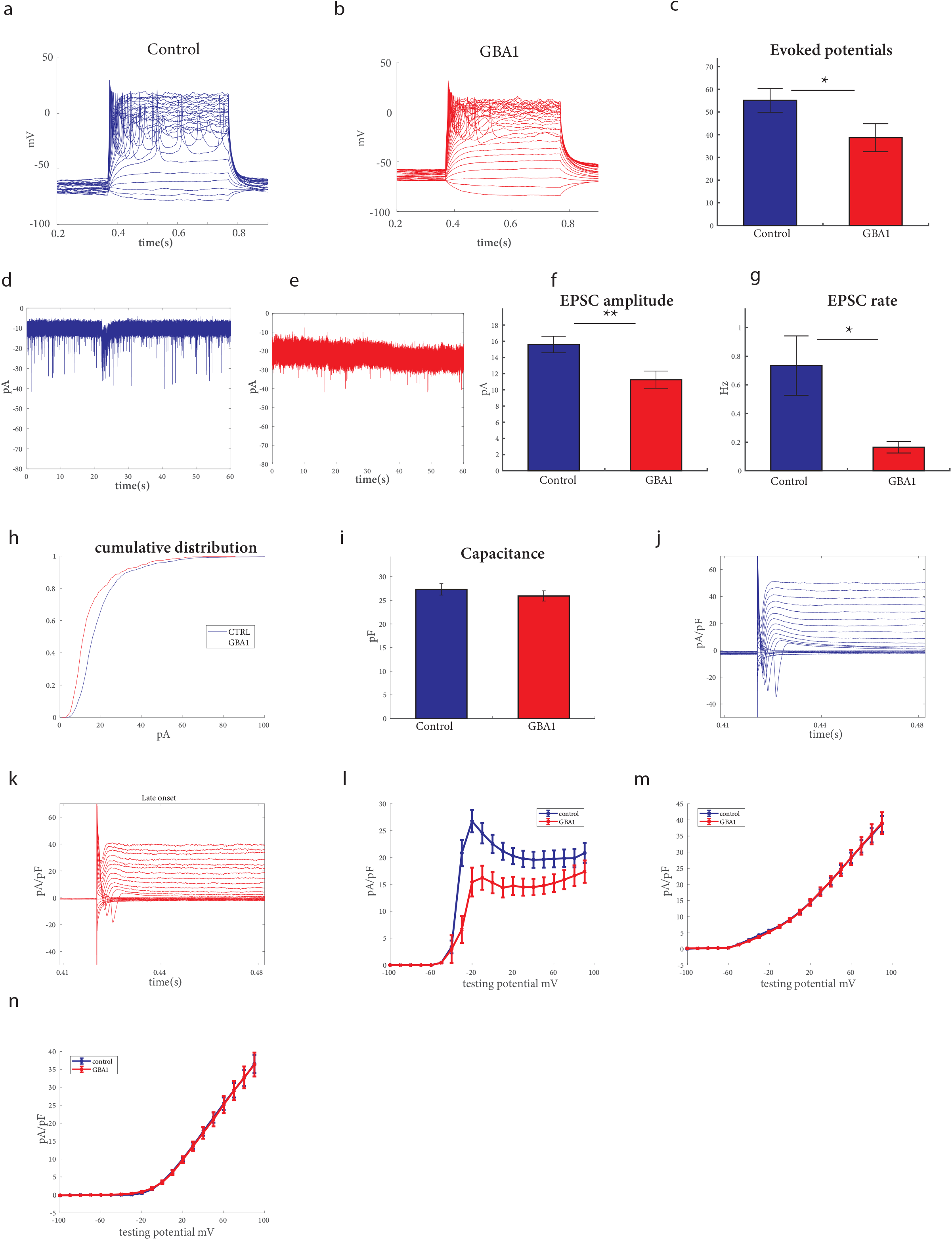
Reduced spontaneous synaptic activity and altered sodium currents in DA neurons derived from GBA1 PD patients compared to healthy controls. iPSC-derived DA neurons from three control individuals and three PD patients with GBA1 mutations were measured by whole-cell patch clamp. (a)-(b) Representative images of evoked potentials of DA neurons derived from GBA1 PD patients and controls, respectively. (c) The total number of evoked action potentials was significantly lower in *GBA1* mutant neurons than in control neurons. (d)-(e) Example recordings of excitatory post-synaptic currents (EPSCs) in control and *GBA1* mutant neurons, respectively. (f) The mean amplitude of EPSCs was significantly reduced in *GBA1* mutant neurons compared to control neurons. (g) The EPSC rate was significantly lower in *GBA1* mutant neurons compared to control neurons. (h) The cumulative distribution of EPSC amplitudes of *GBA1* mutant and control neurons indicates lower EPSC amplitudes in *GBA1* mutant neurons. (i) No significant difference was observed in the capacitance of the *GBA1* and control neurons. (j)-(k) Examples of the recordings of sodium and potassium currents obtained in the voltage-clamp mode from control and *GBA1* mutant neurons, respectively. (l) Our findings indicate a significant decrease in the sodium currents between - 20mV to 0mV in *GBA1* mutant neurons compared to control neurons. (m) No significant differences were observed in the fast potassium currents between 40mV to 80mV between *GBA1* mutant and control neurons. (n) No significant differences were observed in the slow potassium currents between 40mV to 80mV between *GBA1* mutant and control neurons. Asterisks in this and the subsequent figures denote statistical significance as indicated by the following codes: * p<0.05, **p<0.01, *** p<0.001, **** p<0.0001.

Figures 1(j)-(k) present examples of the recordings of sodium and potassium currents obtained in the voltage-clamp mode from control and *GBA1* mutant neurons, respectively. Our findings indicate a significant reduction in the sodium currents between -20 - 0mV in *GBA1* mutant neurons compared to control neurons (Figure 1(l), p=0.002). However, we observed no significant differences in the fast potassium and slow potassium currents between 40-80mV in *GBA1* mutant neurons compared to control neurons (Figures 1(m)-(n), with a one-way ANOVA analysis). Overall, our results suggest that the *GBA1* mutation leads to a reduction in the sodium currents and synaptic activity in DA neurons.

### ECM receptor interaction, Focal-adhesion, and pathways in cancer are amongst the strongest dysregulated pathways in *GBA1* mutant DA neurons

Subsequently, RNA was extracted from DA neurons derived from three *GBA1* mutant PD patients and three control individuals, and gene expression differences were analyzed. Figure 2(a) illustrates DEGs; Specifically, 398 upregulated and 431 downregulated genes were identified in the *GBA1* DA neurons compared to healthy controls (|log2 fold change| > 0.5, FDR < 0.05). The signaling network analysis with the top enriched KEGG pathways of the *GBA1* mutant DA neurons compared to the controls is presented in figure 2(b). The top dysregulated KEGG pathways are shown in figure 2(c) and include the ECM receptor interaction, focal adhesion, and Pathways in cancer. The ECM receptor interaction pathway plays a crucial role in the regulation of synaptic plasticity (72), while focal adhesion proteins maintain the adhesion of brain cells to the ECM, which is crucial for neuronal migration and neurogenesis (73) as well as cell survival (74, 75) and synaptic integrity (77). Additionally, several developmental biological functions were dysregulated in the mutant group compared to the healthy control, as illustrated in Figure 2(d) such as Tissue morphogenesis and Tissue remodeling which involve the organization and restricting of cells and ECM components (78).

**Figure 2:**
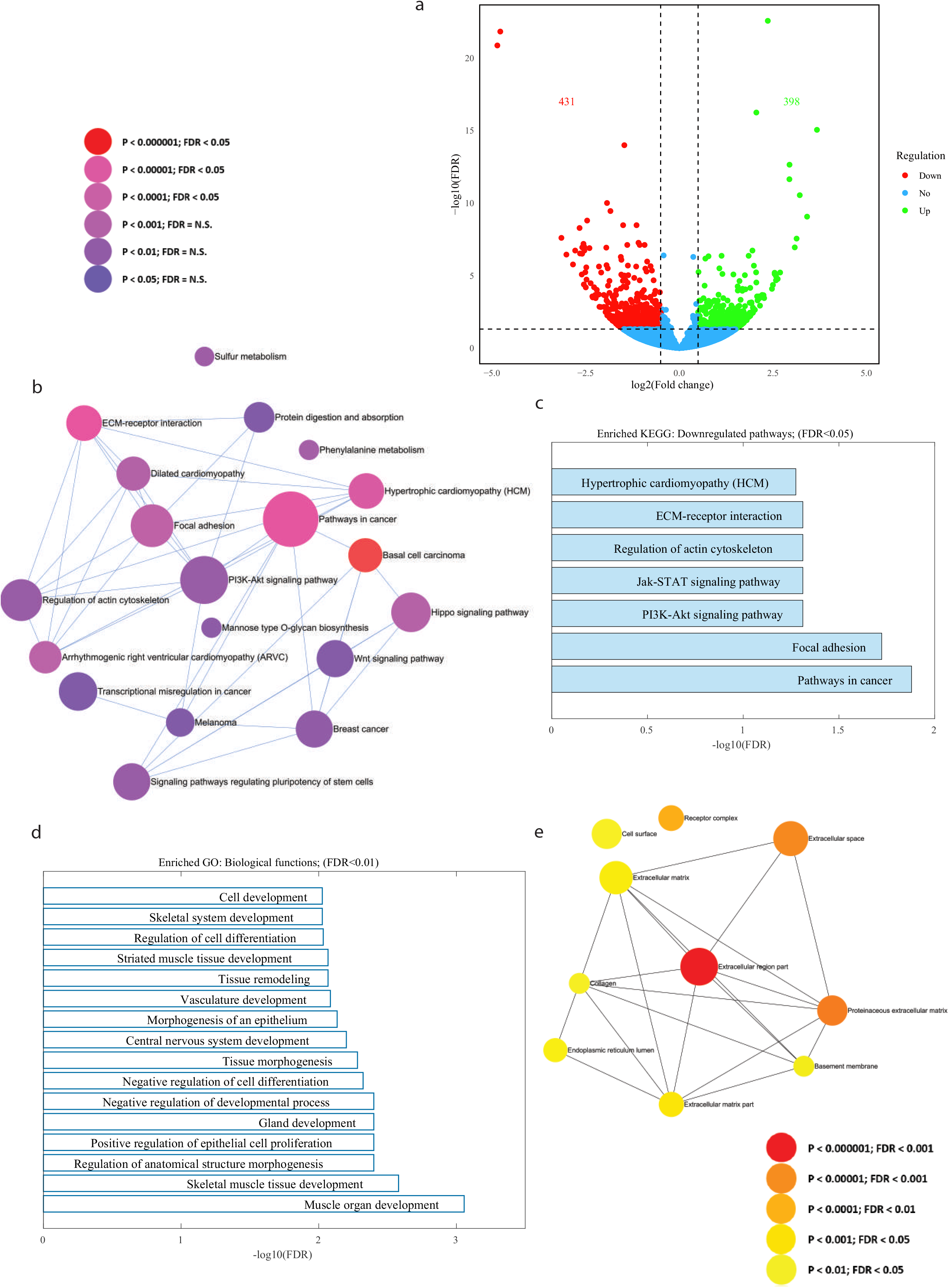
The top dysregulated pathways in GBA1 mutant neurons relate to ECM, focal adhesion, and PI3K-Akt signaling. RNA was extracted from three *GBA1* PD and three healthy control neuronal DA cultures. (a) A volcano plot of DEGs between *GBA1* PD mutant DA neurons and healthy controls (|log2 fold change| > 0.5, FDR < 0.05). (b) Signaling network analysis with the top enriched KEGG pathways in *GBA1* PD mutant DA neurons compared to healthy controls (FDR < 0.05), the size of each node is proportional to the number of DEGs. (c) The top dysregulated (downregulated) KEGG pathways in the *GBA1* PD mutant DA neurons compared to healthy controls (FDR < 0.05). (d) The top dysregulated GO biological functions in the *GBA1* PD mutant DA neurons compared to controls (FDR < 0.01). (e) Signaling network analysis with the top enriched GO Cellular Components in the *GBA1* mutant DA neurons compared to healthy controls (FDR < 0.05).

Furthermore, several extracellular matrix-related cellular components were dysregulated in the mutant group compared to healthy controls, as shown in Figure 2(e). Overall, similar to our previous reports (26, 27), we see that neurons derived from PD patients with different mutations have convergent dysregulated pathways, many of them are related to the ECM and genes that produce adhesion proteins of the cells to the ECM. The PI3K-Akt signaling pathway is also a convergent dysregulated pathway.

### sPD DA neurons exhibit alterations in synaptic activity that are dependent on the age onset of the disease

Using whole-cell patch-clamp, we recorded from a total of 53 control, 27 early-onset sPD neurons, and 16 late-onset sPD neurons derived from 3 control, 3 early-onset sPD, and one patient with a late-onset of the disease. Figure 3 (a)-(c) provides representative traces of evoked potentials in a control neuron, an early-onset sPD neuron and a late-onset sPD neuron. Our findings indicate that the excitability of late-onset sPD neurons is increased compared to control and early-onset sPD neurons. The average excitability is depicted in Figure 2(d) (p=0.49 for control vs. early-onset sPD, p=0.00003 for late-onset sPD vs. control, and p=0.0008 for late-onset sPD vs. early-onset sPD).

**Figure 3:**
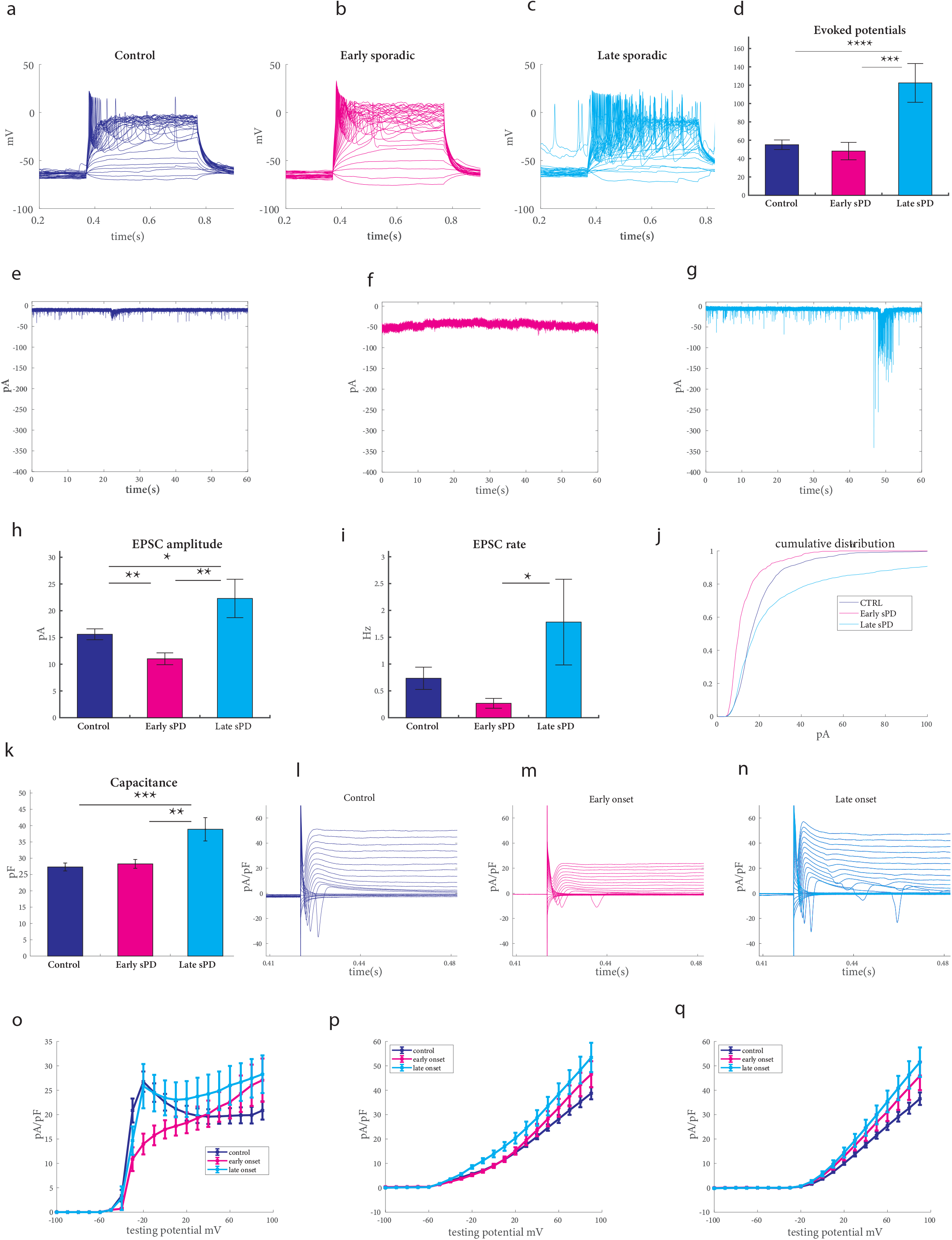
Synaptic dysfunction in DA neurons derived from patients with sPD compared to healthy controls. We recorded from a total of 53 control, 27 early-onset sPD neurons, and 16 late-onset sPD neurons derived from 3 control, 3 early-onset sPD, and one patient with a late-onset of the disease. (a)-(c) Representative traces of evoked action potentials of control, early-onset sPD, and late-onset sPD DA neurons, respectively. (d) Late-onset sPD DA neurons are hyperexcitable compared to control and early-onset sPD neurons as measured by the total evoked action potentials. (e)-(g) Examples recordings of excitatory post-synaptic currents (EPSCs) in control, early-onset sPD, and late-onset sPD DA neurons, respectively. (h) Reduced amplitudes of EPSCs in early-onset sPD and increased in late-onset sPD as compared to the control DA neurons. (i) The rate of EPSCs is increased in late-onset sPD compared to early-onset sPD. (j) A cumulative distribution of EPSC amplitudes in control, early onset sPD, and late-onset sPD neurons. (k) The capacitance is increased in late-onset sPD compared to control and early-onset sPD neurons. (l-n) Examples recordings of the sodium and potassium currents obtained in voltage-clamp mode from control, early-onset sPD, and late-onset sPD neurons. (o) Sodium currents are decreased in early-onset sPD DA neurons compared to controls and late-onset sPD. (p) Fast potassium currents and (q) slow potassium currents are increased in late-onset sPD DA neurons compared to controls.

Our measurements of the synaptic activity revealed significant changes that are dependent on the disease onset age. Figures 2(e)-(g) provide examples of EPSC recordings from a control, early-onset sPD, and late-onset sPD DA neurons measured in a voltage clamp of -60 mV. There was a significant increase in the mean amplitude of EPSCs in late-onset sPD neurons compared to controls (Figure 2(h), p=0.016 for control vs. late-onset sPD neurons, p=0.001 for early-onset sPD vs. late-onset sPD neurons). On the other hand, early-onset sPD neurons exhibited a significant decrease in the EPSC amplitudes compared to control neurons (Figure 2(h), p=0.008). Interestingly, we observed a higher EPSC rate in late-onset sPD neurons compared to control neurons, although this difference was not significant (Figure 2(i), p=0.07). However, a significant difference was noted when we compared the large EPSC rate (see Methods) (Supp. Figure 1(a), p=0.025). In contrast, the EPSC rate was similar in early sPD DA neurons and control neurons, while it was significantly decreased in early-onset sPD neurons compared to late-onset sPD neurons (Figure 2(i), p=0.11 for control vs. early onset sPD neuron, p=0.018 for early vs. late onset sPD). We conducted a comparison of the percentage of synaptically active cells (see Methods) between early-onset sPD and control neurons, revealing a reduction of synaptically active neurons in early-onset sPD (Supp. Figure 1(b), p=0.01). This agrees with our previous report (26) andindicates that the synaptic connectivity is affected and reduced in early-onset sPD (age 45 of disease onset in our previous report), but to a lower degree than most monogenic PD mutations. Figure 2(j) illustrates the cumulative distribution of EPSC amplitudes in sPD and control neurons. The cumulative distribution of EPSC amplitudes in early-onset sPD neurons was left-shifted compared to control neurons, suggesting lower EPSC amplitudes. On the other hand, the cumulative distribution of EPSC amplitudes in late-onset sPD neurons was right-shifted, indicating higher EPSC amplitudes in late-onset sPD neurons. Additionally, we found that the capacitance of late-onset sPD neurons was significantly increased compared to early-onset sPD and control neurons, indicating that cells had a larger surface area in late-onset sPD compared to control and early-onset sPD neurons (Figure 2(k), p=0.0002 for control vs. late-onset sPD, p = 0.002 for early-onset sPD vs. late-onset sPD).

Figures 2(l)-(n) demonstrate examples of the recordings of sodium and potassium currents in voltage-clamp mode from control, early-onset sPD, and late-onset sPD neurons. Our findings reveal no significant differences in the sodium currents between -20-0mV in late-onset sPD neurons compared to control neurons, while the sodium currents were significantly reduced in early-onset sPD neurons compared to control neurons (Figure 2(o), p=0.0038). In contrast, the fast potassium currents were significantly increased in late-onset sPD neurons compared to controls (Figure 2(p), p = 0.027). Also, the slow potassium current was significantly increased in late-onset sPD compared to controls (Figure 2(q), p=0.035). The comparison was performed for currents elicited at 40-80 mV.

### ECM, focal adhesion, and synaptic pathways are dysregulated in sPD DA neurons

RNA extraction was performed from sPD DA neurons (three early and one late-onset patient) as well as three controls. Analysis of gene expression differences between patients and control DA neurons was conducted, and the results are presented in Figure 4. Figure 4(a), left shows the number of DEGs in early-onset sPD compared to healthy controls, 4(a), middle presents the DEGs in late-onset sPD compared to controls, and 4(a), right presents the combined sPD compared to control DEGs (|log2 fold change| > 0.5, FDR < 0.05). The signaling network analysis with the top enriched KEGG pathways in the early-onset sPD DA neurons compared to the healthy controls is presented in Figure 4(b). Some of the top dysregulated pathways are shown in Figure 4(c) and include several synapse-related pathways, ECM-related pathways, focal adhesion, and pathways in cancer. Similarly, the top enriched KEGG signaling pathways in late sPD DA neurons compared to healthy controls are presented in Figure 4(d). Specifically, Hippo signaling pathways and cell cycle are dysregulated (upregulated) in late sPD DA neurons, as shown in Figure 4(e). Additionally, ECM-related pathways and focal adhesion pathways are dysregulated in late sPD (Figure 4(e)). Figure 4(f) presents the top enriched KEGG signaling pathways in the combined sPD DA neurons compared to healthy controls, with several pathways being significantly dysregulated (upregulated), such as the ECM-receptor interaction and protein digestion and absorption pathways, as shown in Figure 4(f)-(g).

**Figure 4:**
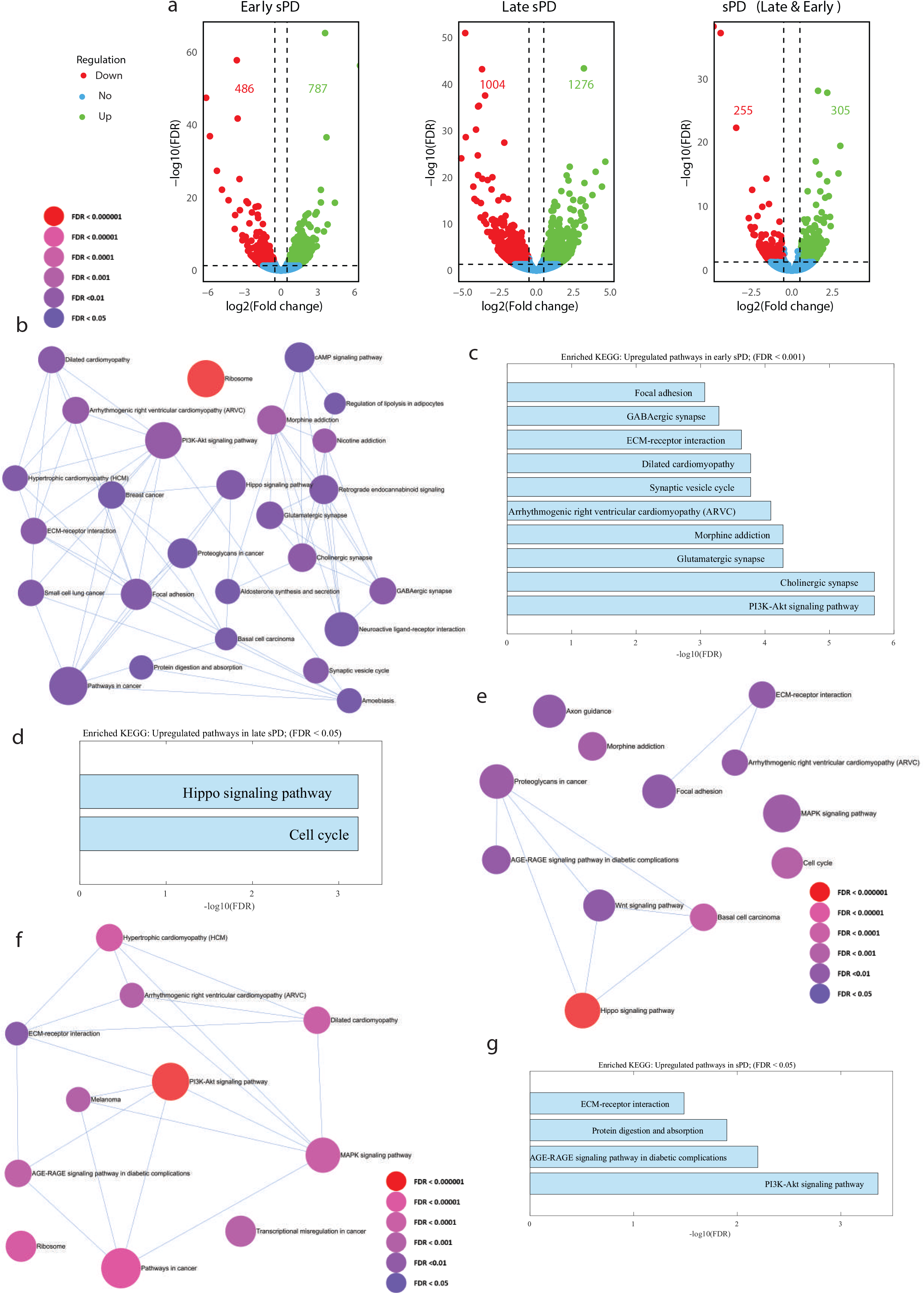
The top dysregulated pathways in sPD DA neurons relate to the Synapse, ECM, focal-adhesion, and PI3K-Akt signaling. RNA was extracted from one early-onset sPD patient, one late-onset sPD patient, and three control individuals’ DA neurons. (a) A volcano plot of DEGs in (a, left) early onset sPD compared to controls. (a, middle) late-onset sPD compared to controls, and (a, right) sPD compared to controls (|log2 fold change| > 0.5, FDR < 0.05). (b) Signaling network analysis with the top enriched KEGG pathways in the early sPD DA neurons compared to healthy controls (FDR < 0.05), the size of each node is proportional to the number of DEGs. (c) The top dysregulated (upregulated) KEGG pathways in the early sPD DA neurons compared to healthy controls (FDR < 0.01). (d) Signaling network analysis with the top enriched KEGG pathways in the late sPD DA neurons compared to healthy controls (FDR < 0.05). (e) The top dysregulated (upregulated) KEGG pathways in the late sPD DA neurons compared to healthy controls (FDR < 0.05). (f) Signaling network analysis with the top enriched KEGG pathways in the combined sPD DA neurons compared to healthy controls (FDR < 0.05). (g) The top dysregulated (upregulated) KEGG pathways in the combined sPD DA neurons compared to healthy controls (FDR < 0.05).

sPD Supplementary analysis revealed additional significant dysregulated cellular functions, including several ECM-related components, cell junction, and neuron projection; Moreover, both sPD early and late onsets showed a dysregulation in several molecular functions and biological processes, including extracellular structure organization, neurogenesis, calcium ion binding, and other related functions, as shown in Supplementary Tables S16-20.

## Discussion

Parkinson’s disease affects up to 3% of people over the age of 80 and also strikes people in their 30s and 40s. Its impact has far-reaching consequences for individuals, families, and society as a whole (75). As a result, extensive research has been conducted, and several PD-causing mutations have been reported; However, the majority of PD cases have no known genetic cause and are classified as sPD cases (4, 9). *GBA1* mutations, which have a significant impact on the pathogenesis of PD, have been reported to have a clear association with the disease (23, 31). In this study, we investigated the intrinsic and network properties of DA neurons derived using iPSC techniques from sPD patients and PD patients who are *GBA1* mutation carriers with no Gaucher disease (heterozygous for the *GBA1* mutation). We additionally analyzed the transcriptome and found some significant changes.

Our findings reveal that the E326K mutation in the *GBA1* gene leads to a reduction in the sodium currents and a decrease in the synaptic activity in DA neurons. We have previously reported alterations in synaptic activity as a shared phenotype of DA neurons derived from PD patients with other mutations (26, 27, 67). It has been demonstrated that reprogramming adult cells into iPSCs removes aging and epigenetic modifications (74, 75). Thus, neurons in our cultures are young neurons. In light of this, the observed phenotype of decreased synaptic activity suggests an early mechanism that is present in the patients’ DA neurons long before the patients experience motor deficits. This finding is consistent with several studies using PD model mice, which suggested that the early phenotype is a decrease in synaptic activity that progressed to motor dysfunction inmany later stages. Thus, monitoring synaptic activity may have a significant impact on the early diagnosis and prediction of PD onset (76-80).

While most PD mutations cause a reduction in synaptic activity (26, 27), PINK1, together with PARK2 mutations, cause an increase in the synaptic activity of DA neurons (27). We report here and previously (26), that early-onset sPD DA neurons’ synaptic activity is similar to most monogenic mutations but with a milder phenotype. However, the late-onset sPD DA neurons had an increase in synaptic activity, as we report here. These observations suggest that any deviation from the normal synaptic activity (even an increase) may have a detrimental effect on neuronal survival. However, the increase in synaptic activity may relate to a disease that occurs at a later age.

Transcriptome analysis and comparison between PD and control DA neurons indicate several intracellular and extracellular pathways that are dysregulated in PD. These include focal adhesion and ECM-receptor interaction which are commonly dysregulated in PD in this study and previous studies (26, 27, 79). These findings suggest a focus on the ECM and its strong involvement in PD. Studies of the ECM composition and structure should be conducted as these are currently under-explored in PD. The synaptic deficits that are also a shared phenotype in PD suggest a temporal correlation between the synaptic impairment and the ECM deficits that needs further investigation in the context of PD. Some of the common intracellular signaling pathways are the PI3K-Akt signaling pathway, cancer-related pathways, Hippo signaling, and more. These intracellular pathways play essential roles in regulating various cellular processes, including cell growth, differentiation, proliferation, and survival. Also, dysregulation of these intracellular pathways has been found relevant in neurodevelopmental disease and neurodegenerative diseases, including Parkinson’s disease (80-85).

The distinct electrophysiological differences between early and late-onset sPD neurons suggest that the underlying molecular mechanisms driving the disease may differ depending on the stage of onset. This notion is further supported by transcriptome analysis, which revealed unique dysregulated pathways in early and late-onset sPD neurons. For example, the upregulation of glutamatergic, GABAergic, and cholinergic synapse pathways, as well as various intracellular signaling pathways in early-onset sPD neurons, might reflect changes in intracellular signaling associated with early disease stages. In contrast, the upregulation of the Hippo signaling pathway and cell cycle in late-onset sPD neurons may be linked to disease progression and cellular changes occurring in the later onset of PD.

Glutamatergic, GABAergic, and cholinergic synapse pathways play essential roles in regulating neurotransmitter signaling within the brain. Glutamate serves as the primary excitatory neurotransmitter, gamma-aminobutyric acid (GABA) functions as the primary inhibitory neurotransmitter, and acetylcholine (ACh) has a modulatory role in controlling neuronal excitability and synaptic plasticity (86, 87). The upregulation of these pathways observed in early-onset sPD neurons might be associated with the decreased EPSC and could represent a compensatory mechanism in response to the reduced synaptic activity. Neurons may attempt to preserve synaptic homeostasis by increasing the expression of genes related to neurotransmitter signaling pathways, thereby counterbalancing the reduction in excitatory input (88). This adaptive response might serve to maintain proper communication between neurons and prevent further deterioration of synaptic function.

In conclusion, this study shows that DA neurons derived from PD patients with GBA1 mutation share synaptic deficits similar to other PD mutations. DA neurons from sPD patients differ in their synaptic abnormalities depending on the age of the onset. Additionally, several shared pathways, such as focal adhesion, ECM-receptor interaction, PI3K-Akt signaling, and more, are commonly dysregulated in monogenic and sporadic forms of the disease. These shared changes in the patients’ DA neurons should now be considered new targets for drug discovery.

## Figure legends

**Supp. Figure 1**

(a) The large EPSC rate was significantly increased in late-onset sPD neurons compared to both control and early-onset sPD. (b) The active cells’ ratio was significantly lower in early-onset sPD neurons compared to control neurons. (c) The input conductance was not different between the control and GBA1 neurons. (d) The input conductance for early-onset sPD neurons was significantly increased compared to control neurons. (e) An example image for DA neurons with Tyrosine Hydroxylase (TH) (green) immunostaining to mark the DA neurons Asterisks in this figure denote statistical significance as indicated by the following codes: * p<0.05, **p<0.01, *** p<0.001, **** p<0.0001.

