## Supplementary figures and images for "Synaptic dysfunction and dysregulation of extracellular matrix-related genes in dopaminergic neurons derived from Parkinson’s disease sporadic patients and with *GBA1* mutations"

### Supplementary figure 1

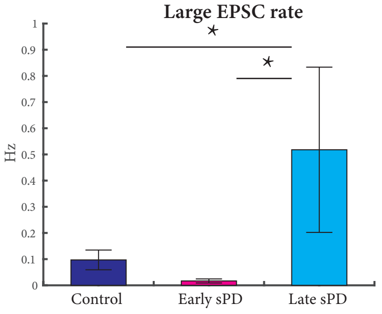

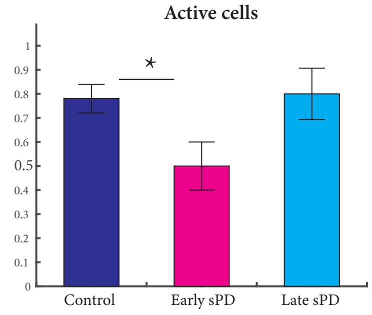

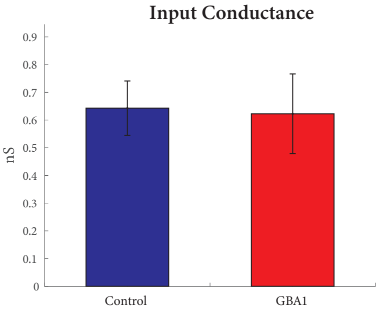

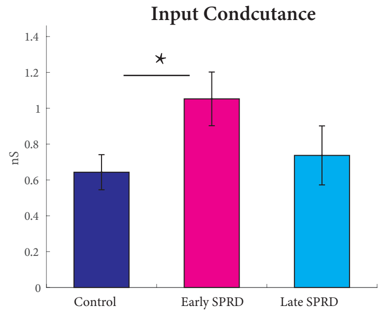

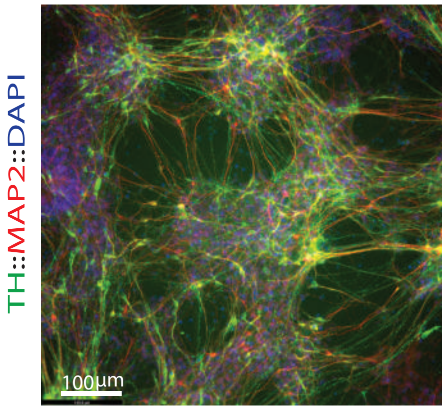
